# Cas12a-Capture: a novel, low-cost, and scalable method for targeted sequencing

**DOI:** 10.1101/2020.11.18.388876

**Authors:** Taylor L. Mighell, Andrew Nishida, Brendan L. O’Connell, Caitlin V. Miller, Sally Grindstaff, Casey A. Thornton, Andrew C. Adey, Daniel Doherty, Brian J. O’Roak

## Abstract

Targeted sequencing remains a valuable technique for clinical and research applications. However, many existing technologies suffer from pervasive GC sequence content bias, high input DNA requirements, and high cost for custom panels. We have developed Cas12a-Capture, a low-cost and highly scalable method for targeted sequencing. The method utilizes preprogramed guide RNAs to direct CRISPR-Cas12a cleavage of double stranded DNA *in vitro* and then takes advantage of the resulting four to five nucleotide overhangs for selective ligation with a custom sequencing adapter. Addition of a second sequencing adapter and enrichment for ligation products generates a targeted sequence library. We first performed a pilot experiment with 7,176 guides targeting 3.5 megabases of DNA. Using these data, we modeled the sequence determinants of Cas12a-Capture efficiency, then designed an optimized set of 11,438 guides targeting 3.0 megabases. The optimized guide set achieves an average 64-fold enrichment of targeted regions with minimal GC bias. Cas12a-Capture variant calls had strong concordance with Illumina Platinum Genome calls, especially for SNVs, which could be improved by applying basic variant quality heuristics. We believe Cas12a-Capture has a wide variety of potential clinical and research applications and is amendable for selective enrichment for any double stranded DNA template or genome.

## Introduction

While advances in sequencing technologies have dramatically reduced the cost and effort required to sequence human genomes, there remains significant clinical and research benefits of targeted sequencing. Restricting genomic interrogation to loci of interest minimizes sequencing costs and analytic labor, reduces data processing and storage, and abrogates ethical issues around returning incidental genetic findings to patients and families. An important application of targeted sequencing is identifying pathogenic variation in Mendelian disorders, which remains a significant clinical challenge as the diagnostic rate for many disorders is only ~50% (1). Likewise, targeted sequencing is a powerful tool for genetic screening, as well as basic research in human or animal genetics. Exome sequencing or hybridization-based gene-panels have been widely used, but these approaches have weaknesses in terms of sequence bias and cost for custom applications. Further, in most cases the non-coding regions of gene bodies are not captured. An easily programmable technology that solved these problems could have major benefits for many clinical and research applications.

CRISPR-Cas systems have emerged as valuable biotechnological tools empowering user-programmed nuclease activity (2, 3). These systems leverage target-specific guide RNAs (gRNAs) to direct Cas9 family endonucleases to specific genomic loci. Several methods have taken advantage of CRISPR-based cutting for capture of genomic regions followed by nanopore (4–9) or Illumina sequencing (9–15). However, some of these rely on laborious size selection steps or require specialized equipment or reagents (8, 13–15). A different set of approaches take advantage of the Cas9-gRNA ribonucleoprotein (RNP) affinity to target DNA by pulling down DNA-bound RNP (10–12). Recently, a method relying on direct adapter ligation to Cas9 cleavage sites, followed by long-read sequencing with nanopore technology has been developed to enable detection of structural variation as well as single nucleotide variants (SNVs) (5–7).

Here, we designed and implemented Cas12a-Capture, a novel, low-cost, and scalable targeted capture technology built around the Cas12a family of enzymes (also known as Cpf1), which cleaves target DNA in a staggered fashion, leaving four to five nucleotide overhangs (16). We reasoned that treating genomic DNA with Cas12a and a pool of gRNAs would result in enrichment of ligatable overhangs specifically at targeted sites. Following ligation of one adapter to the programmed cut sites, a Tn5 transposase is used to add a second adapter at variable distances from cut sites. An enrichment PCR followed by paired end sequencing would then provide targeted sequence data.

To validate the method, we first designed a pilot set and then an optimized set of gRNAs targeting the full gene bodies of genes associated with Joubert Syndrome (JS), a genetically heterogeneous, recessive ciliopathy that manifests with hindbrain malformations (17). The optimized guide set yields strong enrichment with broad coverage of the target region at an affordable price-point. The method also exhibits minimal GC sequence content bias. These characteristics empower sensitive and specific variant calling for whole genes. Cas12a-Capture is currently compatible with Illumina sequencing platforms, but with minor modifications the method could be adapted to other massively parallel sequencing technologies.

## Materials & Methods

### Design of pilot guide set

We obtained RefSeq hg19 genomic coordinates for the 47 genes from UCSC Table Browser as a bed file. Overlapping intervals were merged with Galaxy to obtain a single interval per gene, to which we then padded with 3,000 basepairs upstream and 500 basepairs downstream, in hopes of capturing promoters and 3’ untranslated region sequences. Then, we used FlashFry (18) to find all possible Cas12a target sites (i.e. the presence of “TTTN” PAM) within these target regions and to report the copy number of each potential gRNA target sequence. We filtered out guide target sequences that had copy number greater than one, or that had many similar off target sequences (>25 off targets within 1 edit distance, or >100 off targets within 2 edit distance). We also filtered guides that overlapped a common single nucleotide polymorphism (SNP, minor allele fraction > 0.1%, dbSNP, release 151. Then, for each gene, we defined targets by simply enumerating 500 basepair intervals, and selected the gRNA with cut site closest to the target. This resulted in 7,176 guide sequences. We then designed DNA oligo sequences that contained the following in the 5’ to 3’ direction: dial out PCR priming site, T7 RNA polymerase priming site, crRNA backbone, protospacer sequence, DraI cut-site (“TTTAAA”), and another dial out PCR priming site (Supplementary Table S3). We synthesized these gRNA templates as 99-mers on 12,000-feature oligo chips (CustomArray).

### Guide amplification and *in vitro* transcription of pilot guide set

We used PCR to amplify the gRNA templates from the oligo pool using dialout primers. Reactions contained 1x KAPA HiFi Hotstart Readymix, 10 ng of template, 0.5 μM primers, and 1x SYBR Green. Reactions were pulled upon completing exponential amplification, which occurred at 19-22 cycles. Agarose gel electrophoresis confirmed bands of 99 basepairs. We purified reactions with NucleoSpin PCR cleanup columns (Machery Nagel). Then, we treated purified products with DraI restriction enzyme in order to remove the priming site downstream of the gRNA sequence. Reactions contained 500 ng of PCR product, 40 units of DraI (New England Biolabs), and 1x CutSmart buffer. Incubation was done at 37°C and proceeded overnight. Reactions were cleaned up with NucleoSpin PCR cleanup columns, and complete digestion was confirmed with agarose gel electrophoresis.

We used MEGAscript T7 Transcription Kit (Thermo Fisher Scientific) to generate gRNAs from the templates. Reactions contained ~60-130 ng DNA (depending on recovery from previous step), and were incubated at 37°C overnight. Following incubation, reactions were treated with TURBO DNase and incubated at 37°C for 15 minutes. Then, RNA Clean & Concentrator (Zymo Research) columns were used to purify RNA. We quantified RNA with Qubit RNA Broad Range Assay (Thermo Fisher Scientific) and diluted to 10 μM.

### Cas12a-Capture workflow

For a detailed protocol, see Supplementary Methods. Briefly, genomic DNA is treated with phosphatase to enzymatically remove the terminal phosphates from genomic DNA molecules. Then, genomic DNA is treated with gRNA-complexed Cas12a, which creates overhangs specifically at targeted sites. Custom i5 adapters that contain complementary overhangs, a unique molecular identifier (UMI), and 5’ biotin modification are added with T4 ligase. Then, the i7 adapter is added through Tn5 tagmentation. A streptavidin-mediated pulldown step purifies those molecules that have an i5 adapter (excluding the molecules with only i7 adapters), and on-bead PCR (followed by size selection/purification as necessary) generates ready-to-sequence libraries. All libraries were sequenced in paired-end mode on the Illumina NextSeq500 platform with Mid Output 150 cycle v2.5 kits. Cycles were allocated as follows: 35 cycles for read 1, 10 cycles for index 1, 6 or 10 cycles for index 2 (depending on the presence of unrelated multiplexed libraries), and 113 or 118 cycles for read 2. More sequencing cycles were allocated to read 2 since the read 1 start sites are determined by the Cas12a cut site.

### Sequencing data processing and analysis

Our custom adapter contains a six nucleotide unique molecular identifier (UMI) in place of the i5 index. The first step of our informatics pipeline is appending the sequence from the i5 index read to the end of the read name line of both read 1 and 2 fastq files with a custom python script. This is done for compatibility with UMI-tools (19). Next, adapters are trimmed with cutadapt (20) and paired end reads are aligned to the hg19 reference genome with BWA-MEM (21). Following paired end read alignment, duplicates are removed with UMI-tools dedup.

### Modeling sequence determinants of guide performance

We estimated the performance of guides by the number of sequencing reads that aligned to the predicted cut site. Namely, a read was assigned to a guide if the first base of the read was within the 16^th^ to 26^th^ position downstream of a guide’s PAM. An additional pseudocount read was added to all guide counts, enabling log transformation of all read counts, which we used as the dependent variable. We then collected 667 sequence-based features as in previous work modeling Cas12a *in vivo* activity(22). Four bases upstream of the PAM and six bases downstream of the protospacer were considered. Position-specific nucleotides and dinucleotides were included (excluding the first three positions of the PAM, which are fixed as “T”), as well as two features relating to GC content: the GC imbalance of the protospacer (i.e. how far the actual GC content was from 50%), and the GC content of the predicted overhang (positions 26-30). Additionally, we included the estimated minimum free energy of the RNA molecule(23).

Feature selection was done with the elastic net procedure, implemented in scikit-learn version 0.19.0. We found optimal hyperparameters with cross validation (ElasticNetCV) on 90% of the data (6,447 guides). This procedure resulted in 287 features with non-zero coefficients. To further eliminate inconsequential features, we trained ordinary least squares linear regression models with increasing numbers of features (rank ordered by elastic net coefficient absolute value) and made predictions on the 10% (729) fully withheld guides. Prediction performance did not substantially improve once the top ~100 features were added (Supplementary Figure S2). Therefore, we fit a final ordinary least squares linear regression model to all available data (training and test), with the 100 selected features, which we then used to make predictions for the optimized guide set.

### Design of optimized guide set

We used the same procedure as for the pilot guide set for obtaining padded genomic coordinates, identifying all possible Cas12a target sites, and excluded potential guides with copy number >1 and overlapping SNPs >0.1% allele frequency (dbSNP build 153). We used a restricted list of 34 genes (3.1 megabases), representing high confidence JS risk genes. We also implemented a more sophisticated procedure for picking guides. First, we designed two guide sets, one targeting the forward genomic strand and one targeting the reverse genomic strand, such that consecutive guides alternated orientation. After picking a guide, we defined the next target as 250 basepairs downstream of the predicted cut site. We established a set of criteria, prioritizing high-scoring guides, guides most proximal to the target, and guides with a low number of predicted off target sites. Predicted off target sites for each guide were found by enumerating all possible single nucleotide deletions from the guide sequence and finding perfect matches for these in the genome. If there were no guides of the correct orientation fulfilling the criteria and within 250 basepairs of the target, we broadened our search to guides in the opposite orientation. If there were still no suitable guides, we moved on without choosing a guide. Once this process had been completed for all genes, we identified all “gaps” (i.e. no guides present) of greater than 600 basepairs. We reasoned that flanking the gaps with guides in the optimal orientation (i.e. forward guides upstream and reverse guides downstream of the gap) would maximize our ability to obtain coverage in the gap regions. Initially, 283 gaps representing 264,401 bases were identified. Of these, 140 of these regions already had high scoring guides in the correct orientation flanking the >600 basepairs region, which we considered resolved. This left 143 regions (135,414 basepairs) without flanking guides. If correctly oriented guides were present within 100 basepairs of the gap, regardless of predicted performance, we additionally picked those guides. This scheme fully resolved 43 of these gaps (35,055 basepairs, i.e. picked two new guides that flanked the gap in the desired orientation). A gap was considered partially resolved if we found one flanking guide (86 gaps, 89,935 basepairs) and unresolved if no flanking guides were found (14 gaps, 10,424 basepairs). We picked a total of 11,438 guides for the optimized set, and guides were synthesized as two oPools at the picomole per oligo scale (Integrated DNA Technologies, Supplementary Table S7).

### *In vitro* transcription of optimized guide set

MEGAscript T7 Transcription Kit was used to generate gRNAs directly from the single-stranded templates. We scaled up the recommended reaction volumes five-fold, and added 50 picomoles of oPool template as well as 50 picomoles of T7 promoter. Reactions were incubated at 37°C overnight. Following incubation, reactions were treated with TURBO DNase and incubated at 37°C for 15 minutes. Then, RNA Clean & Concentrator (Zymo Research) columns were used to purify RNA. We quantified RNA with Qubit RNA Broad Range Assay (Thermo Fisher Scientific) and diluted to 10 μM.

### Variant Calling

Base quality scores were recalibrated with BaseRecalibrator from GATK v4.1.2.0. HaplotypeCaller from GATK was run for calling variants using a minimum base quality of 20. Distributions of the metrics from the called variants were created classifying variants as true positives, false positives, and false negatives using variant calls from the confident regions in the NA12878 Platinum Genome (24). Key metrics which separated these classifications, quality by depth (QD) and allele balance (AB), were chosen for F1-score analysis to optimize separate threshold cutoffs for SNVs and indels. Variants were filtered to have an AB < 0.80 and QD > 2.0 for SNVs and an AB < 0.80 and QD > 8.0 for INDELs. Final classifications of variant calls and calculations of precision and recall were made using hap.py v0.3.12-2-g9d128a9. Variant classification was constrained to confident regions from Platinum Genomes and stratified by repeat regions taken from RepeatMasker and low complexity BED files from the Global Alliance for Genomics and Health (GA4GH) Benchmarking Team and the Genome in a Bottle Consortium (25, 26).

## Results

### Overview of the Cas12a-Capture method

The key intuition of the approach is that Cas12a-mediated genomic fragmentation, mediated by a pool of targeted gRNAs, should result in enrichment of ligatable overhanging ends at targeted loci. Moreover, Cas12a cleavage can occur completely *in vitro* on naked DNA. Specific gRNAs can be generated in bulk at low cost by synthesizing pools of DNA oligonucleotides containing the gRNA sequence as well as the T7 RNA polymerase priming site. *In vitro* transcription is then used to generate pools of functional gRNAs. In order to reduce spurious ligation events, genomic DNA can be enzymatically dephosphorylated prior to incubation with the Cas12a-gRNA RNP (Figure 1). Cas12a cleavage results in a 5’ overhang of four to five nucleotides. Therefore, we designed custom, biotinylated adapters containing the Illumina i5 flow cell and priming sequences, as well as overhangs of four or five degenerate nucleotides (Supplementary Table S1). Following ligation of the i5 adapter, tagmentation with Tn5 transposase adds the i7 sequencing adapter. Finally, to enrich for molecules with a ligated i5 adapter (and deplete molecules with two i7 adapters), a streptavidin-mediated pulldown is performed, followed by PCR directly on the streptavidin beads (Figure 1, full protocol can be found in Supplementary Methods).

**Figure 1.**
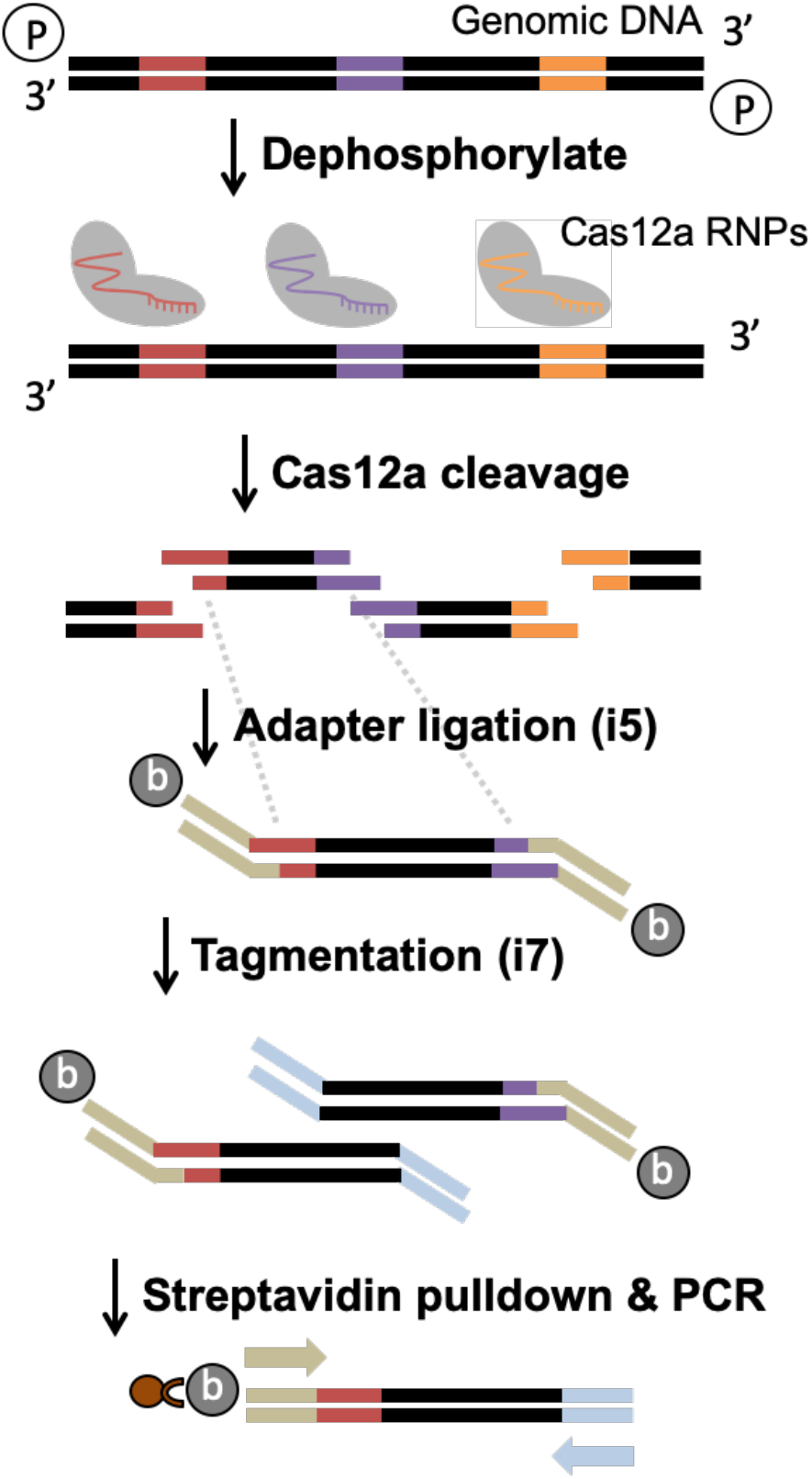
Schematic outlining the Cas12a-Capture protocol. Genomic DNA (black bars) is dephosphorylated and then treated with Cas12a as well as a pool of gRNAs that cleave target sites (colored regions on genomic DNA) to leave overhangs. A custom biotinylated adapter (beige bars) with degenerate overhangs is ligated to the cleaved molecules, then the other adapter (light blue) is added with Tn5 tagmentation. Finally, a streptavidin pulldown isolates library molecules which are amplified by on-bead PCR (beige and light blue arrows are primers).

### Design and performance of the pilot guide set

To validate the method and learn the sequence determinants of capture efficiency, a pilot set of guides was designed targeting 47 known and candidate genes associated with JS (Supplementary Table S2), representing 3.5 megabases of DNA. The only design criteria were the presence of a “TTTN” protospacer adjacent motif (PAM), and filters for guides overlapping SNPs or with predicted off target cut sites (Materials and Methods). The design tiled 7,176 guides at 500 base-pair intervals. DNA oligonucleotides encoding the T7 RNA polymerase promoter, *Acidaminococcus sp.BV3L6* (*As*) Cas12a constant loop region, and target specific protospacer region were produced with array-based synthesis (Supplementary Table S3). Subsequent *in vitro* transcription produced mature gRNAs. Combined paired-end sequencing data from several Cas12a-Capture libraries prepared from the well-studied CEPH/Hapmap sample NA12878 resulted in 5.9% of reads on target, corresponding to a 52.4-fold enrichment. While this enrichment was encouraging, we sought to understand the source of off target reads. As the primary error modality of array synthesis is single base deletions, we generated a predicted off target list by aggregating all sites in the genome at which gRNAs with a single base deletion aligned (495,299 sites). We observed 12.7% of sequencing reads aligned to these predicted off target sites, which is significantly more than aligned to the same number of size-matched random genomic intervals (1.75%, p < 0.01, Chi-squared test). Since Cas12a cleavage results in symmetrical 5’ overhangs, we expected that approximately equal numbers of reads would result from ligation to both overhangs. However, this was not the case: 56% of guides had greater than 10 times more reads aligning to the enzyme-distal overhang (Supplementary Figure S1). This bias may be due to Cas12a remaining bound to the enzyme proximal fragment (27) and sterically inhibiting ligation, though treatment with sodium dodecyl sulfate after cleavage did not reduce the bias (data not shown).

Inspection of the read alignments revealed accumulation of the first read at programmed cut sites, consistent with ligation of the i5 adapter directly to the cut site overhang. We found for the first read of 92.6% of on target read pairs began within 5 bases of a predicted guide cut site. Additionally, we observed that the starting position of the first read corresponds to the expected cut sites of Cas12a (16) (i.e. after the 18^th^ and 23^rd^ bases downstream of the PAM, Supplementary Figure S1). In contrast, the second read was scattered across the inter-guide interval, consistent with this adapter being appended by semi-random tagmentation (Supplementary Figure S1).

### Modeling Cas12a-Capture performance

Comparing the performance of capture across the full guide set revealed a thousand-fold difference between the best and worst performing guides; however, 49.3% of guides performed within one log10 difference (Figure 2A). We reasoned that we should be able to use the pilot data as a training set to model the sequence determinants of Cas12a-Capture performance. Toward this end, we collated 667 sequence-based features, representing position-specific nucleotides and dinucleotides, GC content, and gRNA folding (Supplementary Table S4). We used the number of reads assigned to each guide as a proxy for the performance of that guide. We modeled Cas12a-Capture performance using linear regression and implemented elastic net regularization to assign hyperparameters and feature coefficients (22). Hyperparameters were chosen with nested cross-validation, and we tested the resulting model on fully withheld data. The predicted and observed scores were highly correlated (Pearson r = 0.79, Figure 2C). Overall, 287 features were assigned non-zero coefficients. Based on a plateau in predictive performance, we used the top 100 features in the final model (Supplementary Figure S2). Consistent with previous work, a thymine at the fourth position of the PAM is strongly disfavored (16). Other important features were related to GC content. GC imbalance of the guide was strongly disfavored. High GC content at the overhang positively related to performance, likely due to increased ligation efficiency (Figure 2B). Inspection of contributions from single position-specific nucleotides suggests the most important positions are within the seed and overhang regions (Figure 2D).

**Figure 2.**
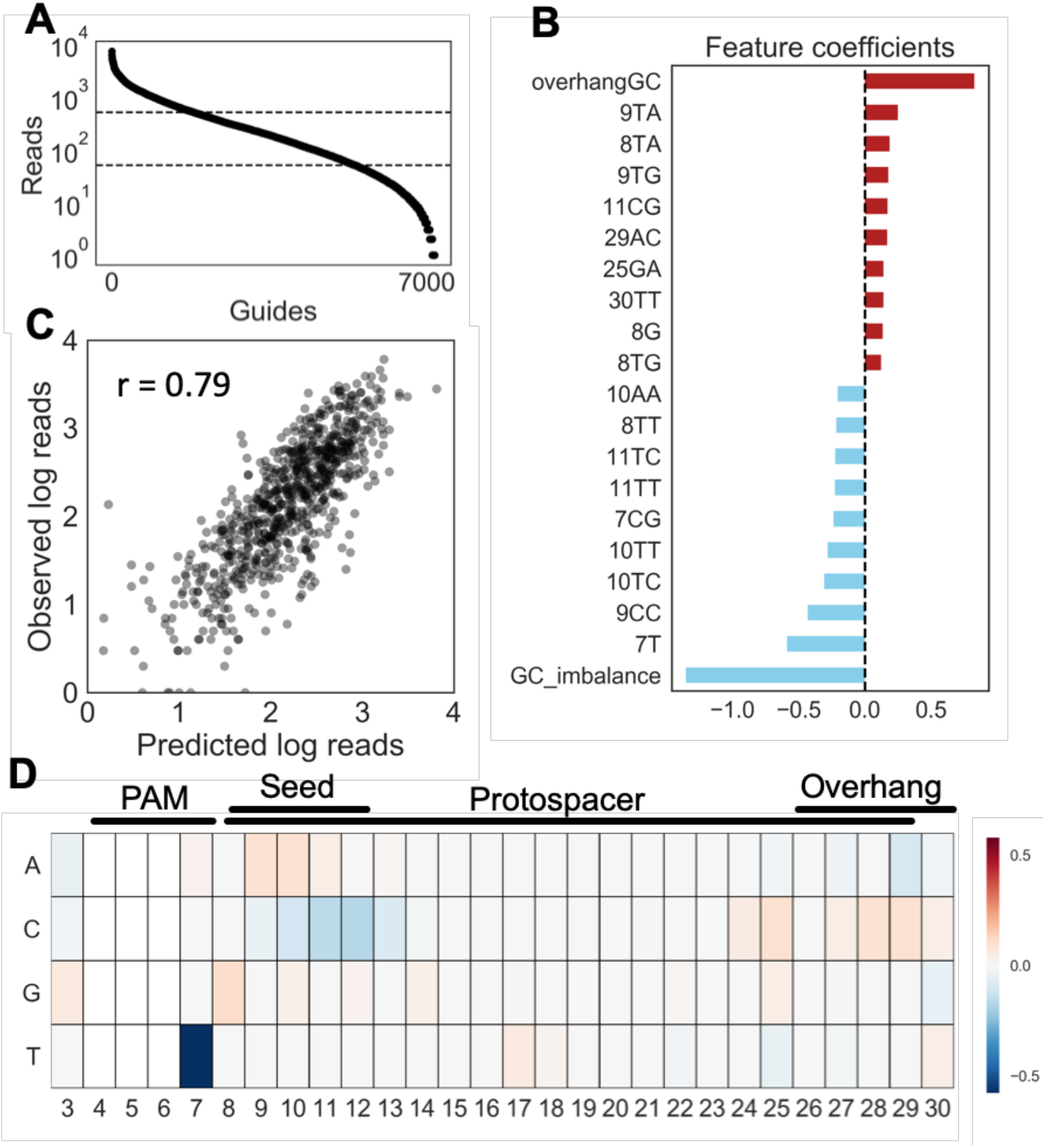
Modeling the sequence determinants of Cas12a-Capture performance. (**A**) Read uniformity for guides in the pilot experiment. Dashed lines indicate a log10 window within which 49.3% of guides performed. (**B**) The twenty features in the linear regression model with the largest positive and negative coefficients. (**C**) Cross validation of the linear regression model on fully withheld test data. Pearson r = 0.79. (**D**) Feature coefficients of individual position-specific nucleotides in the DNA target.

### Design and testing of optimized guide set

We used a combination of predicted performance scores from our model, optimal spacing, and number of predicted off target cut sites to generate optimized guide sets for 34 high confidence JS risk genes using a higher fidelity column-based synthesis platform (Materials & Methods, Supplementary Table S6). Due to the observed biased capture efficiency from enzyme distal versus proximal fragments, we designed two interleaved pools, one targeting the forward genomic strand and the other targeting the reverse genomic strand (Supplementary Table S7). Following an initial guide selection step, we identified gaps (>600 basepairs between subsequent guides) and, when possible, picked additional guides flanking the gaps in an attempt to capture the gap sequence (Materials & Methods).

Captures across two experiments, two replicates each, with the optimized guides achieved an average enrichment of 64-fold (6.3% of reads on target) using NA12878 genomic DNA. On average, 80.7% of reads aligned to the genome and 97.6% of these were unique (Supplementary Table S8). Guide uniformity improved modestly compared to the naïve guide set with 54.0% of guides within one log10 difference (Figure 3A). While cutting at predicted off target sites was present, it made up a relatively smaller fraction of reads compared to the pilot guides (5.5% vs. 12.7% of reads). Observed guide performance correlated with predictions (Pearson r = 0.38, Supplementary Figure S3), but this correlation was lower than the cross-validation results.

**Figure 3.**
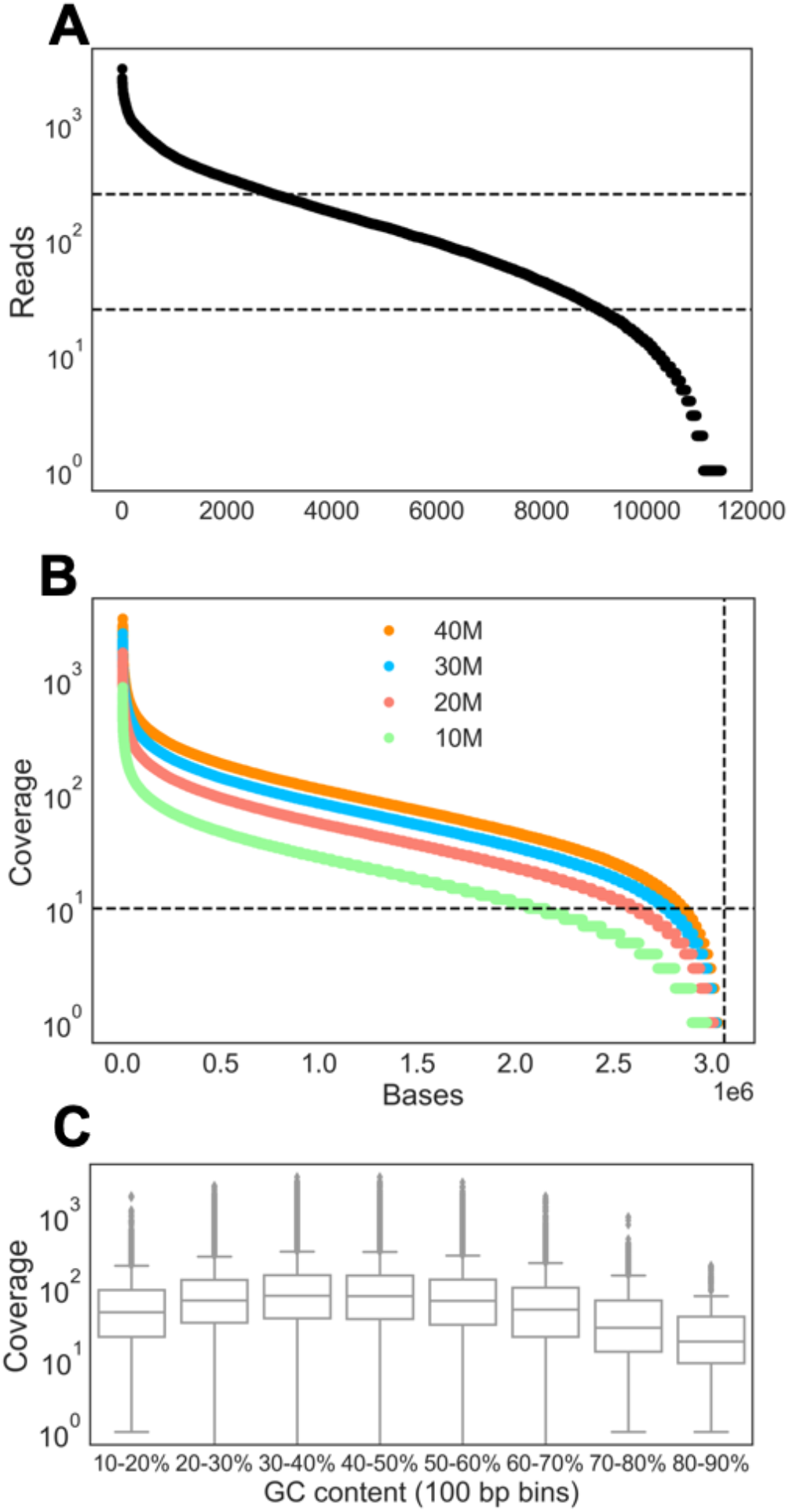
Performance of the optimized guide set. (**A**) Read uniformity for guides in the optimized experiment. Dashed lines indicate a log10 window within which 54.0% of guides performed. (**B**) Per-base read coverage across the full target with downsampled datasets. (**C**) Boxplots showing coverage of bases within different 100 basepair GC content bins.

Combining data across different replicates of NA12878 captures, we generated a high depth dataset and examined coverage of the target region at different levels of downsampling. With 20 million read pairs, 84.4% of bases in the target region are covered by at least 10 reads, and increasing to 30 or 40 million read pairs increases coverage to 90.0% or 92.8%, respectively (Figure 3B). Considering only those bases outside of repetitive elements (as defined by Repeat Masker), 20 million read pairs cover 86.7% of bases with at least 10 reads, and at 40 million read pairs 94.6% of bases are covered by at least 10 reads (Supplementary Figure S3). Coverage for the protein coding sequences of the target genes was similar to overall coverage (Supplementary Table S9).

Examining the potential design gap regions that required selecting suboptimal guides (4.4% of total target), we found similar coverage in the 40 million read set for gaps resolved with two correct orientation flanking guides (88% at least 10 reads, Supplementary Table S10). Unresolved and partially resolved gaps (i.e. only one flanking guide) had lower target coverage (64% and 54% at least 10 reads, respectively). We next examined GC content coverage bias. 100 basepair bins with extremely low (10-20%) or high (80-90%) GC content have median coverage of 46 and 18, respectively, while the 40-50% bin has median coverage of 78 (Figure 3C).

Finally, we performed SNV and insertion/deletion (indel) calling with the 40 million read pair dataset. We compared our calls to NA12878 calls from the Illumina Platinum Genomes confident regions (24, 26) and explored the suitability for various quality metric heuristics to distinguish accurate from inaccurate calls (Supplementary Figure S4, Materials & Methods). Performance of SNV calling without heuristics was high with precision and recall of 0.94 and 0.90, respectively (Supplementary Table S11). Performance improves using heuristic filtering (Materials & Methods), precision and recall of 0.97 and 0.89, respectively (Table 1). We next compared the performance of the filtered calls outside of or within different repeat defined regions. Unsurprisingly, we found lower SNV performance within repeat regions.

**Table 1.** Variant calling performance using filtering heuristics

| **<u>SNVs</u>** | **Precision** | **Recall** | **Truth Total<sup>a</sup>** | **TP** | **FP** | **FN (<10 reads)** |
| --- | --- | --- | --- | --- | --- | --- |
| **All** | **0.97** | **0.89** | 2080 | 1861 | 66 | 219 (182) |
| **Excluding Repeat Masker** | **0.98** | **0.92** | 965 | 892 | 20 | 73 (43) |
| **Within Repeat Masker** | **0.95** | **0.87** | 1115 | 969 | 46 | 146 (139) |
| **Excluding Homopolymers and Tandem Repeats<sup>b</sup>** | **0.97** | **0.91** | 1908 | 1732 | 57 | 176 (142) |
| **Within Homopolymers and Tandem Repeats** | **0.93** | **0.76** | 172 | 129 | 9 | 43 (40) |
| **<u>Indels</u>** |  |  |  |  |  |  |
| **All** | **0.86** | **0.60** | 394 | 238 | 39 | 156 (105) |
| **Excluding Repeat Masker** | **0.82** | **0.55** | 188 | 103 | 22 | 85 (37) |
| **Within Repeat Masker** | **0.93** | **0.76** | 206 | 135 | 17 | 71 (68) |
| **Excluding Homopolymers and Tandem Repeats** | **0.91** | **0.89** | 136 | 121 | 12 | 15 (7) |
| **Within Homopolymers and Tandem Repeats** | **0.93** | **0.76** | 258 | 117 | 27 | 141 (98) |

<sup>a</sup>Eberle *et al.*, 2017.
<sup>b</sup>taken from https://github.com/genome-in-a-bottle/genome-stratifications and see Krusche *et al.*, 2019.
Abbreviations: TP-true positive, FP-false positive.

For indel calls, overall performance without heuristics was poor with precision and recall of 0.50 and 0.65, respectively. However, performance was substantially improved with heuristic filtering to an indel precision and recall of 0.86 and 0.60, respectively (Table 1). Unexpectedly, performance within the Repeat Masker defined regions was improved, while the non-Repeat Masker region performance was similar to the overall target. In contrast, regions of the target outside of homopolymers and tandem repeats showed substantially improved metrics for precision (0.91) and recall (0.89), suggesting most of the indel calling challenges are driven by issues related to these complex regions.

## Discussion

In this study, we demonstrate and evaluate Cas12a-Capture, a novel method for targeted enrichment of regions of interest for sequencing. This method takes advantage of the unique cleavage pattern of Cas12a to introduce ligatable overhangs specifically at regions of interest. In a pilot experiment, we measured capture efficiency for 7,126 gRNAs, chosen without any sequence-specific criteria, and used these data to model the sequence determinants of capture efficiency with high accuracy. Evaluation of an optimized guide set designed to capture the whole bodies of 34 genes revealed 10x coverage of ~84-93% of all target bases with 20-40 million paired end reads.

Major strengths of the method are affordability and flexibility. Synthesis of gRNA-encoding DNA oligos is quick and inexpensive (~$0.01/base), and standard *in vitro* transcription kits can generate thousands of reactions worth of gRNA from picomolar scale DNA template, effectively making the oligos a one-time cost that amortizes well for moderately sized cohorts. For ongoing studies, gRNAs targeting different genes could easily be added to an existing pool. Further, there is no requirement for specialized equipment, and the protocol can be completed in a single day. Though not implemented here, the method is compatible with liquid handling robots that could dramatically increase throughput. GC sequence content bias of Cas12a-Capture compares favorably with hybridization-based approaches (28), and represents a novel approach for inexpensive and sensitive small variant discovery in whole genes.

Our method is the latest in an expanding suite of CRISPR-based targeted sequencing approaches. Some methods seek to characterize repeat expansions (4, 13) or fusion genes (5, 6), while others are aimed at enriching one or a few loci (8, 11, 14, 15). The published approaches that are most comparable to ours are CRISPR-Cap (10) and nCATS (7). In CRISPR-Cap, biotinylated Cas9-RNP enables pulldown by remaining bound to cleaved targets. At a comparable 100 nanograms input DNA, the authors report that CRISPR-Cap enriches a 164 kilobase target 117.9-fold using Illumina sequencing. This enrichment was achieved with guides tiled every 20 basepairs, requiring substantially more guides than our design. For nCATS, Cas9 cleavage at target sequences is used to generate ligation compatible ends in otherwise dephosphorylated genomic DNA similarly to Cas12a-Capture, followed by Oxford Nanopore sequencing. This method achieved a range of 366.5 to 807.0-fold enrichment of a 177 kilobase target with an input of 3 μg input DNA from NA12878. This amount of input may be difficult to obtain, particularly for patient or archived samples. An achievement that sets our method apart from other CRISPR-based targeted sequencing approaches is the high multiplexing of the target space, here 3×10^6^ bases. Thus, while our current enrichment metrics are less than some Cas9-based methods, we are able to target ~20-fold more regions. Additionally, we develop a model for accurately predicting guide efficiency which has not been done by other groups. A weakness of this approach with current Illumina sequencing platforms is the inability to span longer repeat elements, such as LINEs that are common within gene bodies, or detect other more complex structural variants. However, our method would be compatible with longer-read nanopore sequencing with modifications to the ligation adaptor.

The main design constraints for Cas12a-Capture are the dependence on a PAM site and a target sequence with high Cas12a cutting efficiency. We were able to identity high scoring guides in our model for 93.6% of our design target, suggesting that most of the human genome will be accessible to this method. Moreover, for regions with poor scoring guides, we were still able to recover >50% of the target sequence. Efforts are under way, though, to find Cas12a variants with more flexible PAM requirements (29), increased cutting efficiency (30), and reduced off target cutting (31). Additionally, while we achieve strong enrichment of targeted regions, the majority of sequencing reads originate from off target loci. We attribute a substantial fraction of off target reads to synthesis errors in the gRNA encoding DNA oligos. We show that higher fidelity column-based synthesis reduces the percent of putative off target cutting, and oligo purification schemes that eliminate deletion errors would also likely be beneficial. An improved understanding of the origin of the rest of the off target reads could lead to substantial improvement in the percent of on target reads.

While our modeling improved capture performance, the observed guide performance showed weaker overall correlation compared to the cross-validation results. This is likely due to the optimized guides falling within a narrower range of expected performance compared to the cross-validation. For example, when only considering guides above the 2.0 threshold used for picking optimized guides, the cross-validation results in reduced correlation (Pearson r = 0.61). Additionally, the pilot guides were subjected to PCR amplification and restriction enzyme digestion steps prior to *in vitro* transcription while the optimized guides were not. These additional steps could introduce biases that are not present for the optimized guide set. Finally, we observe an imbalance in the capture efficiency for enzyme proximal versus enzyme distal cleavage product. Cas12a (27) and Cas9 are known to remain bound to cleavage products and this forms the basis of the CRISPR-Cap (10) method. A possible explanation for our results is some Cas12a remaining bound to cleavage products and sterically interfering with subsequent ligation, despite heat inactivation of Cas12a and column cleanup of the cleavage reaction prior to ligation.

We foresee broad utility of the method in Mendelian disease genetics, where it can be valuable to sequence the full bodies of custom lists of genes. However, the method is compatible with *in vitro* targeting of any DNA genome or double-stranded DNA library (e.g., cDNA), and could also be useful in basic research, diagnostics, or agriculture.

## Acknowledgements

We would like to thank Jack Weidrick and Ryan Mulqueen for helpful discussions. We would like to thank Andy Fields for help preparing Tn5 transposase.

## Author Contributions

TLM, ACA, and BJO developed the initial outline of the Cas12a-Capture method. BJO, TLM, and DD designed the study. BJO and DD supervised the study. BLO, DD, ACA, and BJO helped design and interpret experiments. TLM, CAT, SG, and CVM performed the experiments. TLM and AN analyzed the data and produced the figures and tables. TLM and BJO drafted and edited the manuscript with input from all authors.

