## Supplementary Methods and Figures S1-S4 for "Cas12a-Capture: a novel, low-cost, and scalable method for targeted sequencing"

### Cas12a-Capture Protocol

#### Reagents

- Shrimp alkaline phosphatase (rSAP, New England BioLabs, Cat. M0371)
- Phosphate buffered saline (PBS, Thermo Fisher, Cat. 10010023)
- 10x Cas9 reaction buffer (recipe below, from IDT)
- Alt-R A.s. Cas12a (Cpf1) V3 (IDT, Cat. 1081068)
- NucleoSpin Gel and PCR Clean-Up (Takara, Cat. 740609)
- Custom i5 adapter (Supplemental Table 1)
- T4 DNA Ligase (New England BioLabs, Cat. M2622)
- T4 DNA Ligase Buffer (New England BioLabs, Cat. B0202)
- TAPS (Sigma Aldrich, Cat. T5130)
- Potassium Acetate (Sigma Aldrich, Cat. P1190)
- Magnesium Acetate (Sigma Aldrich, Cat. M5661)
- DMF (Sigma Aldrich, Cat. D4551)
- Loaded Tn5 transposase (Picelli et al., 2014. Genome Research)
- Sodium Dodecyl Sulfate (SDS, Sigma Aldrich, Cat. L3771)
- Dynabeads MyOne Streptavidin C1 (Thermo Fisher, Cat. 65002)
- NaCl (Fisher, Cat. M-11624)
- Tris (Fisher, Cat. T1503)
- LiCl (Sigma Aldrich, Cat. L9650)
- EDTA (Sigma Aldrich, Cat. E9884)
- Tween-20 (Sigma Aldrich, Cat. P1379)
- KAPA HiFi Hotstart ReadyMix (Roche, Cat. KK2602)
- Nextera i7 indexed primers (Supplemental Table 1)
- Sera-Mag Select SPRI beads (GE Healthcare, 29343045)

#### Equipment

- Magnetic tube rack
- DNA Engine Tetrad Thermal Cycler (BioRad, or other thermal cycler)
- Thermomixer (Thermo Fisher, Cat. 5382000015, or other heat block)
- Illumina sequencing instrument

#### Protocol

1. **Prepare transposome**
    a. Hybridize i7_adapter_top and i7_adapter_bottom (each at 100 µM).
        i. Combine 8 µL each of top and bottom adapters with 52.4 µL of 2x Tn5 dilution buffer (20 mM HEPES-KOH, 200 mM NaCl, 25% glycerol, 0.2% Triton-X, 2 mM DTT).
        ii. Incubate at 95°C for 5 minutes.
        iii. Cool to 20°C at 2.5°C per minute.
    b. Adjust NaCl concentration of Tn5.
        i. Combine 1152 µL of 16 µM Tn5 (see Picelli *et al.*, 2014, Genome Research) with 144 µL of 5M NaCl.
    c. Load transposase

    i. Combine equal volumes of hybridized i7 adapters and NaCl adjusted Tn5
    ii. Incubate at 25°C for 1 hour.
    iii. Store at -20 for no more than 8 months.

2. **Dephosphorylate genomic DNA**
    a. Prepare 10x Cas9 reaction buffer (can be done beforehand)
        i. 200 mM HEPES, 1μM NaCl, 50 mM $MgCl_2$, 1 mM EDTA, pH 6.5 @ 25°C
    b. Combine 100 ng genomic DNA (quantified with Qubit fluorometer) with 2 μL of 10x Cas9 buffer, 2 μL rSAP, and water to total volume of 20 μL.
    c. Incubate at 37°C for 30 minutes.
    d. Incubate at 65°C for 5 minutes.

3. **CRISPR cleavage of genomic DNA**
    a. Dilute Cas12a to 1 μM with PBS.
    b. Combine 7.5 μL water, 1.5 μL of 10x Cas9 reaction buffer, 2 μL of 10 μM gRNA, and 4 μL of 1 μM Cas12a.
    c. Incubate at room temperature for 10 minutes.
    d. Combine the gRNA / Cas12a mixture with the reaction from step 1.
    e. Incubate at 37°C for 30 minutes.
    f. Incubate at 65°C for 10 minutes.
    g. Purify DNA with Nucleospin columns; elute in 20 μL of buffer NE.

4. **Ligate i5 adapter**
    a. Anneal adapter oligos (can be done beforehand and frozen).
        i. Combine 10 μL i5_adapter_top, 5 μL i5_adapter_bottom_4N, 5 μL i5_adapter_bottom_5N, and 80 μL TE.
        ii. Heat to 95°C in thermal cycler, then, cool at a rate of 0.1°C/second until reaching 10°C.
    b. To the eluate from step 2, add 0.5 μL water, 2.5 μL T4 DNA Ligase Buffer, 1 μL T4 DNA Ligase, and 1 μL of 10 μM annealed i5 adapter.
    c. Incubate at 25°C for 30 minutes.
    d. Incubate at 65°C for 10 minutes.

5. **Tagment DNA**
    a. Make 1 mL fresh 4x TAPS buffer: 132 μL of 1M TAPS, 52.8 μL of 5M potassium acetate, 40 μL of 1M magnesium acetate, 640 μL of 100% DMF, 135.2 μL water
    b. To the reaction from the previous step, add 12.5 μL 4x TAPS buffer, 11.5 μL water, and 1 μL 8 μM loaded indexed Tn5 transposase.
    c. Incubate at 55°C for 5 minutes.
    d. Transfer to ice.
    e. Add 5 μL of 2% SDS.
    f. Incubate at room temperature for 5 minutes

6. **Streptavidin magnetic bead pulldown**
    a. Prepare following buffers (can be done beforehand):
        i. LWB: 10 mM Tris-Cl pH 8.0, 1M LiCl, 1mM EDTA, 0.05% Tween-20, in water.
        ii. NWB: 10 mM Tris-Cl pH 8.0, 1M NaCl, 1mM EDTA, 0.05% Tween-20, in water.
        iii. TWB: 10 mM Tris-Cl pH 8.0, 0.5mM EDTA, 0.05% Tween-20, in water.
        iv. 2x NTB: 10 mM Tris-Cl pH 8.0, 2M NaCl, 1mM EDTA, in water.

   b. Warm Dynabeads MyOne Streptavidin C1 beads to room temperature for 30 minutes.
   c. For each sample, transfer 5 µL beads to PCR tube in a magnetic rack.
   d. Concentrate beads (until supernatant is clear) and remove supernatant.
   e. Wash with 200 µL TWB, concentrate, remove supernatant.
   f. Resuspend beads in 110 µL 2x NTB.
   g. Add samples to beads; shake for 30 minutes at 1,000 rpm at room temp.
   h. Wash 1x with 200 µL LWB. Concentrate and remove supernatant.
   i. Wash 2x with 200 µL NWB. Concentrate and remove supernatant.
   j. Wash 2x with 200 µL TWB. Concentrate and remove supernatant.
   k. Resuspend in PCR mix: 12.5 µL KAPA HiFi HotStart ReadyMix, 0.5 µM custom i5 primer, 0.5 µM indexed Nextera i7 primer, water to 25 µL. Ensure that beads are dispersed (i.e. have not settled to the bottom of tube).

7. **On-bead PCR and cleanup**
   a. Thermal cycle: 72°°C for 3 minutes, 95°°C for 30 seconds, repeat 15 total times: 98°°C for 20 seconds, 60°°C for 15 seconds, 72°°C for 40 seconds.
   b. Concentrate Dynabeads and transfer supernatant to a new tube.
   c. Cleanup PCR with Sera-Mag Select SPRI beads, at a 0.8x beads to sample ratio. Check size distribution with preferred method.
   d. Multiple rounds of SPRI bead cleanup may be necessary. If libraries become too dilute after multiple cleanups, a “reconditioning PCR” may be performed, in which the libraries are re-amplified (<= ~5 cycles) with primers that bind the flow-cell binding portion of the sequencing adapters.
8. **Perform paired end sequencing on an Illumina instrument**
   a. Example read lengths for 150 cycle kitRead lengths should be:
      i. Read1: 35 cycles
      ii. Index1: 10 cycles
      iii. Index2: 6 cycles
      iv. Read2: 117 cycles

**Supplementary Figures**

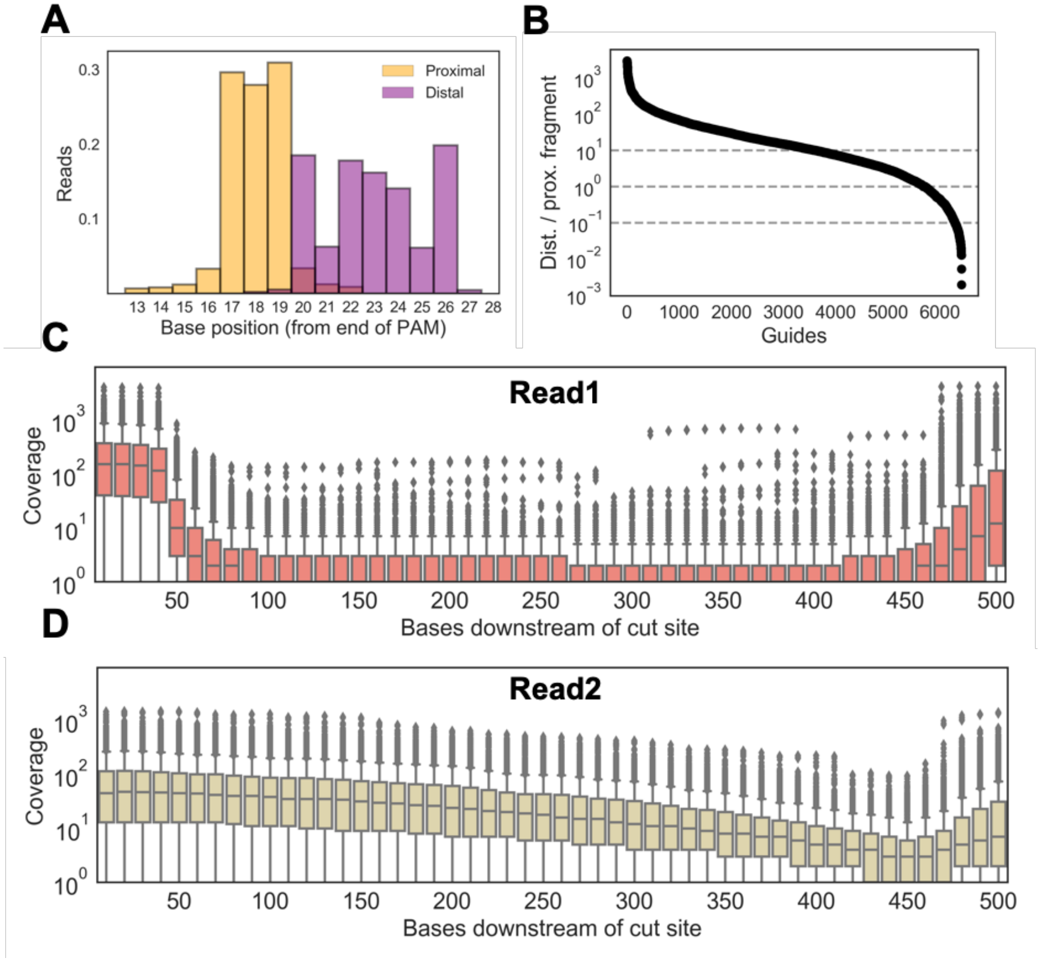


**Figure S1.** Characteristics of CRISPR-Capture.
(**A**) Histogram of position of first base of read 1 in relation to the end of PAM (i.e. the start of the protospacer). Reads originating from the Cas12a proximal and distal molecules are colored differently.
(**B**) Ratio of Cas12a distal to proximal reads for all guides, rank ordered by magnitude of ratio.
(**C**) Boxplots showing coverage of bases, from read 1, as a function of distance downstream from nearest cut site.
(**D**) Boxplots showing coverage of bases, from read 2, as a function of distance downstream from nearest cut site.

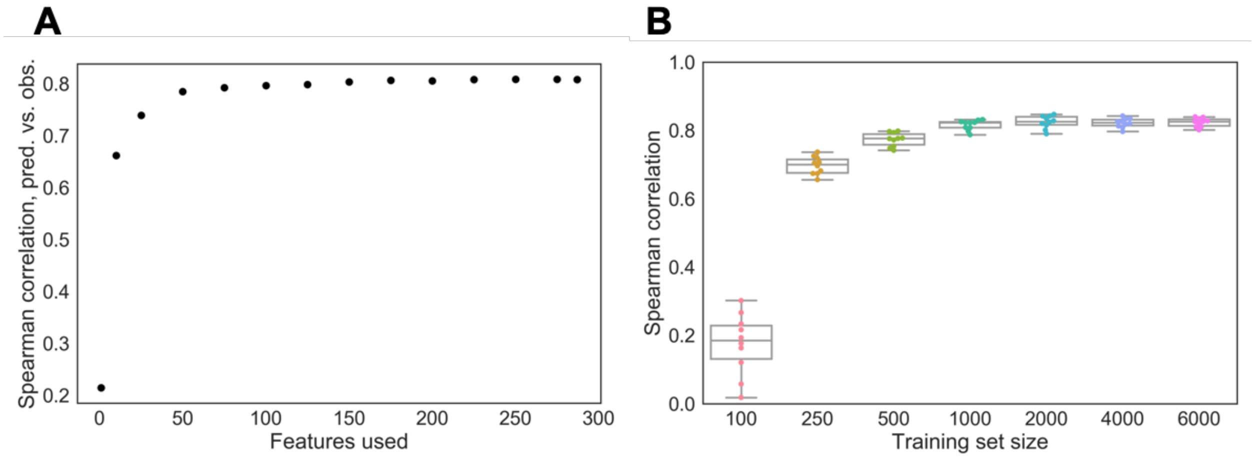


**Figure S2.** Modeling sequence determinants of CRISPR-Capture performance.

(**A**) Models were iteratively trained with more features, successively adding features with the highest absolute value coefficient.

(**B**) Boxplots showing Spearman correlation for models trained with varying numbers of gRNAs from the pilot probe set.

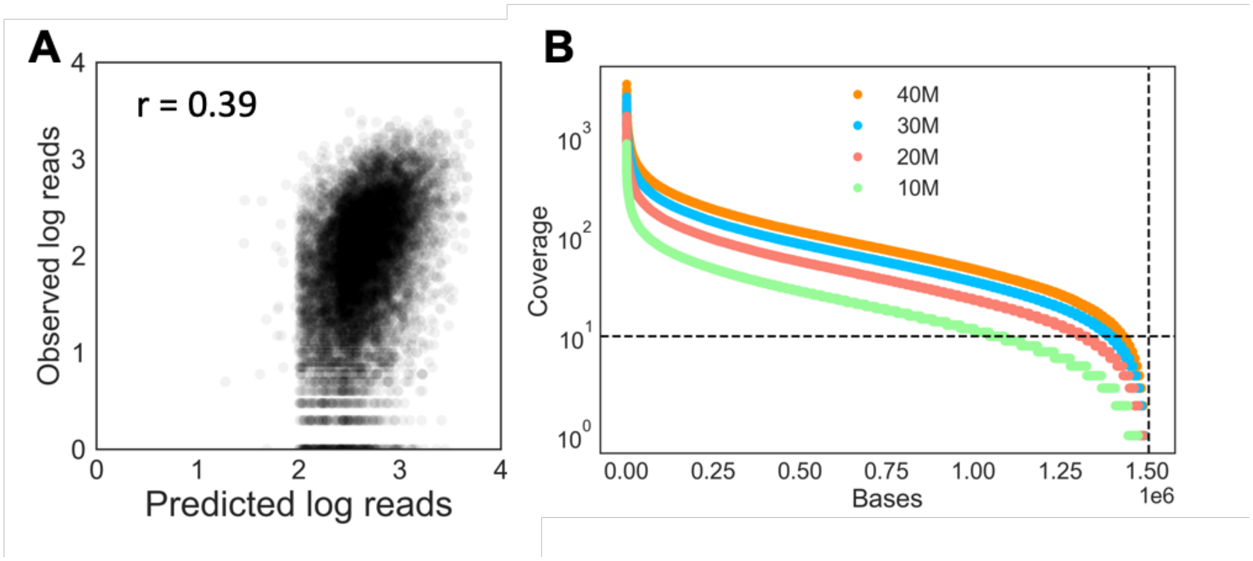


**Figure S3.** Performance of optimized guide set.
(**A**) Predicted versus observed performance (as defined by assigned reads) for the optimized guide set. Pearson r = 0.39. Both axes log10 scale.
(**B**) Coverage uniformity for all bases outside of repeats (as defined by Repeat Masker) for various downsampled datasets.

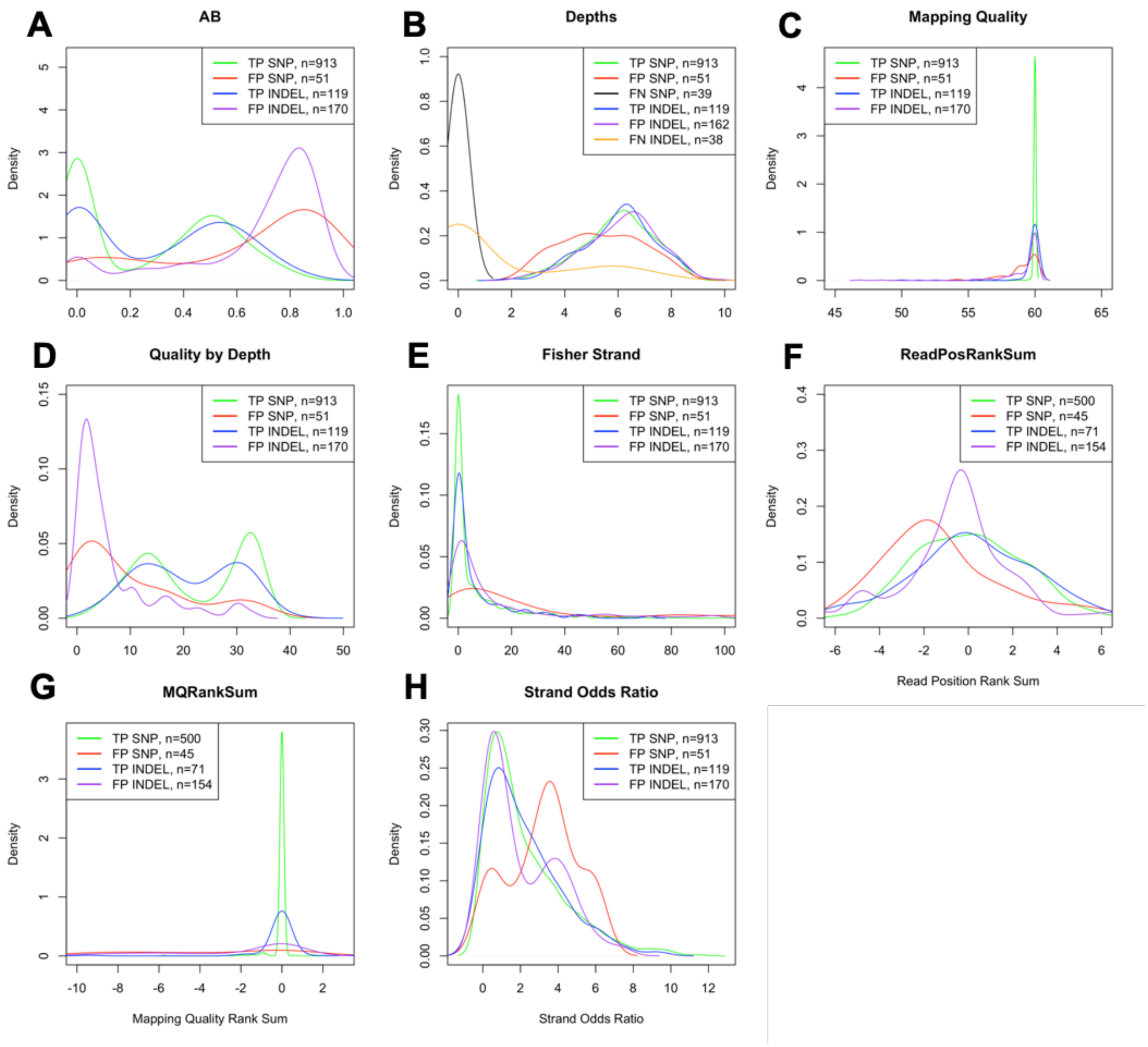


**Figure S4.** Variant calling metrics across entire target region (including repeats) with 40 million read pairs.
(**A**) Allele balance (AB) for true or false positive SNVs or indels.
(**B**) Read depth (DP) for true or false positive SNVs or indels. False negative SNPs and indels are also included.
(**C**) Mapping quality for true or false positive SNVs or indels.
(**D**) Quality by depth (QP) for true or false positive SNVs or indels.
(**E**) Strand bias estimated with Fisher exact test for true or false positive SNVs or indels.
(**F**) Read position bias estimated with rank sum test for true or false positive SNVs or indels.
(**G**) Rank sum test for mapping qualities (MQ) of true or false positive SNVs or indels.
(**H**) Strand bias estimated by the symmetric odds ratio test for true or false positive SNVs or indels.
